# Intracranial pressure elevation alters CSF clearance pathways

**DOI:** 10.1101/760645

**Authors:** Vegard Vinje, Anders Eklund, Kent-Andre Mardal, Marie E. Rognes, Karen-Helene Støverud

## Abstract

**Background:** Infusion testing is a common procedure to determine whether shunting will be beneficial in patients with normal pressure hydrocephalus. The method has a well-developed theoretical foundation and corresponding mathematical models that describe the CSF circulation from the choroid plexus to the arachnoid granulations. Here, we investigate to what extent the proposed glymphatic or paravascular pathway (or similar pathways) modifies the results of the traditional mathematical models.

**Methods:** We used a two-compartment model consisting of the subarachnoid space and the paravascular spaces. For the arachnoid granulations, the cribriform plate, capillaries and paravascular spaces, resistances were calculated and used to estimate flow before and during an infusion test. Next, pressure in the subarachnoid space and paravascular spaces were computed. Finally, different variations to the model were tested to evaluate the sensitivity of selected parameters.

**Results:** At baseline, we found a very small paravascular flow directed into the subarachnoid space, while 60% of the fluid left through the arachnoid granulations and 40% left through the cribriform plate. However, during the infusion, paravascular flow reversed and 25% of the fluid left through these spaces, while 60% went through the arachnoid granulations and only 15% through the cribriform plate.

**Conclusions:** The relative distribution of CSF flow to different clearance pathways depends on intracranial pressure (ICP), with the arachnoid granulations as the main contributor to outflow. As such, ICP increase is an important factor that should be addressed when determining the pathways of injected substances in the subarachnoid space.

## Background

Infusion testing is a standard procedure to assess whether patients with normal pressure hydrocephalus (a type of dementia) would benefit from shunt surgery. During infusion of artificial cerebrospinal fluid (CSF), intracranial pressure (ICP) is monitored, and a CSF outflow resistance (*R*_out_) is calculated. Typically, a constant infusion rate of 1.5 ml/min results in an ICP pressure increase by around 10–25 mmHg, and the calculated *R*_out_ parameter is commonly used as a supplementary parameter in the selection of patients for shunt surgery [41]. The procedure has a well developed theoretical foundation as well as corresponding mathematical models (see [42] for an overview). The main outflow route is assumed to be the arachnoid granulations (AG) [40] in accordance with the traditional view of the third circulation where CSF is produced in the choroid plexus and absorbed through AG as proposed by Cushing in 1925 [10].

More recently, an alternative CSF circulation has been proposed – the glymphatic circulation. Here, paravascular spaces, extensions of the Virchow–Robin spaces, play an active role in brain-wide CSF circulation in conduits that runs in parallel with the vasculature. The purpose of this circulation is to clear solutes from deep inside the brain, thus taking the role of the lymphatic system within the central nervous system wich is abscent of lymphatic vessels. Therefore, this waste clearance system has been named the glymphatic system [33], where the “g” indicates that glial cells play an important role. Glymphatic dysfunction has been hypothesized to contribute to development in neurodegenerative disorders, traumatic brain injury and stroke [33]. In the glymphatic circulation, CSF moves through the subarachnoid space (SAS) along arteries and dives into the brain along arterial paravascular spaces (PVS). The glymphatic pathway continues through AQP-4 channels, through extracellular spaces (ECS) and eventually reaching the venous PVS.

The main bulk of evidence for the glymphatic pathway has been established via in-vivo rodent experiments. In these experiments, tracers are typically infused in the CSF in rodents at a rate of 0.34–2 *μ*L/min, with a resulting pressure increase of 0.1–2.5 mmHg [31, 5, 46]. Thus, such tracer experiments may in fact be viewed as infusion tests. This potential link, between infusion tests and the glymphatic system, has not yet been explored.

Recently, the resistance of the glymphatic system under normal conditions was estimated by Faghih & Sharp [19]. They concluded that the glymphatic circulation was unlikely, as the high resistance of the pathway would prevent sufficient flow. In their model, the resistance of the paraarterial tree was relatively low before reaching the precapillary level where the PVS was set to 100 nm in accordance with a study of Bedussi et al. [5]. The narrow PVS at the capillary level effectively blocked the circulation. However, other studies suggest flow within the paravascular spaces at the level of capillaries [65, 25]. Furthermore, it has been argued that fixation, which was used by Bedussi et al. [5], shrinks the PVS [46]. As such, the resistance of the PVS at the capillary level should be further investigated and compared to the high resistance to flow through the ECS of the brain parenchyma [29, 62, 54].

Several other possible CSF outflow pathways have been proposed. In sheep it has been reported that outflow through the cribriform plate plays a major role in CSF absorption, whereas the importance of the AG is unclear [59]. Flow through the cribriform plate has been suggested to dominate the paravascular flow route when total CSF efflux is large [38]. In addition, the Bulat-Klarica-Orešković hypothesis states that production and absorption is present everywhere in the CSF system, especially over the capillary wall due to its large surface area [9]. Other CSF outflow routes have also been proposed [50, 27], however, a quantification of the fluid distribution and interplay between each outflow pathway is yet to be properly addressed. In addition, resistance of flow from the paraarterial space through the ECS and/or along capillaries in the setting of infusion tests (i.e. under temporarily elevated pressure) has not yet been investigated. We note that lumbar intrathecal contrast delivery during infusion to assess glymphatic function in humans was proposed in Yang et al. [70]. Further, as suggested by Ma et al. [38] increased flow, and a possible change in ICP, may alter the distribution of CSF to different outflow pathways. In addition, if the glymphatic circulation is a main outflow route for CSF, the outflow resistance *R*_out_ is a direct measure of glymphatic dysfunction, which in turn has been linked to neurodegenerative disorders [33].

On this background, the aim of this work was to quantify different CSF outflow routes in the setting of an infusion test. To do so, we first gathered and summarized available resistances of the more probable outflow pathways including the AG, the cribriform plate, the arterial and venous PVS, and CSF drainage/filtration over the capillary wall. We next estimated resistances in the missing segments, i.e. the capillary gaps and the ECS. With these resistances, we extended a well-established mathematical infusion model to include additional pathways and then assessed the relative importance of the different outflow routes at baseline ICP and during infusion of fluid into the CSF system. We modeled an infusion test to explore potential changes in CSF outflow routes with the rising pressure. The relative importance of each outflow route was found to change with increasing ICP, although clearance through AG was dominant both at baseline and (elevated) plateau ICP. At baseline, flow in PVS was relatively stagnant, with an average velocity of 0.04 *μ*m/s from the PVS into the SAS, but reversed with an average velocity of 1.97 *μ*m/s from the SAS into PVS at plateau ICP.

## Methods

### Exit routes from the SAS

Mathematical models of the infusion test have usually assumed the AG to be the main exit route from the SAS [43, 12, 14, 40, 16]. Our purpose here was to investigate and quantify the plausibility of other routes, in particular in light of the proposed glymphatic system [30, 33]. It should be noted that this is our interpretation of the glymphatic system on the macroscale, the original hypothesis include both micro- and macroscopic processes. More precisely, we assumed three exit routes for CSF leaving the SAS, namely the AG entering the dural sinus, the cribriform plate (crib) entering into lymphatic vessels, and PVS [27]. CSF entering the paravascular spaces may enter the brain parenchyma, possibly through AQP-4 channels, [30], or continue along the PVS to reach the PVS at the precapillary level. From the precapillaries, we assumed three further possible pathways: absorption by the capillaries (cap) to leave the system through the bloodstream, flow into the ECS, or flow along small gaps around capillaries [65, 25, 49]. Fluid from both the ECS and capillary gaps eventually return to the paravenous spaces. From the paravenous spaces, the glymphatic circulation suggests a re-entry into the SAS [33]. However, such a circulation is not compatible with a pressure driven flow through the brain parenchyma, since that would require a pressure difference between the input and output of the paravascular space. To allow for circulation, we here introduce a hypothetical compartment, with an associated pressure, downstream from the paravenous compartment. This additional compartment models further transport along spaces potentially anatomically distinct from the SAS such as e.g. paravenous spaces surrounded by pial sleeves [71]. Although compelling evidence for such a route is lacking, a direct contact between paravenous spaces and lymphatic vessels has been hypothesized by some investigators [56]. It should also be noted that a direct route to cervical lymphatics was also envisioned in the paper first proposing the term “glymphatic pathway” [30].

Figure 1 shows the full model, as an extension of the model often used in the literature assuming outflow through the AG only [14,16]. Throughout, we will compare results from the full model, as shown in Figure 1, to a previously established and clinically accepted model (Figure 2), referred to as the reference model. This reference model is based on one lumped outflow resistance *R*_out_ and a pressure-dependent compliance. Several full model modifications will be described and considered.

**Figure 1:**
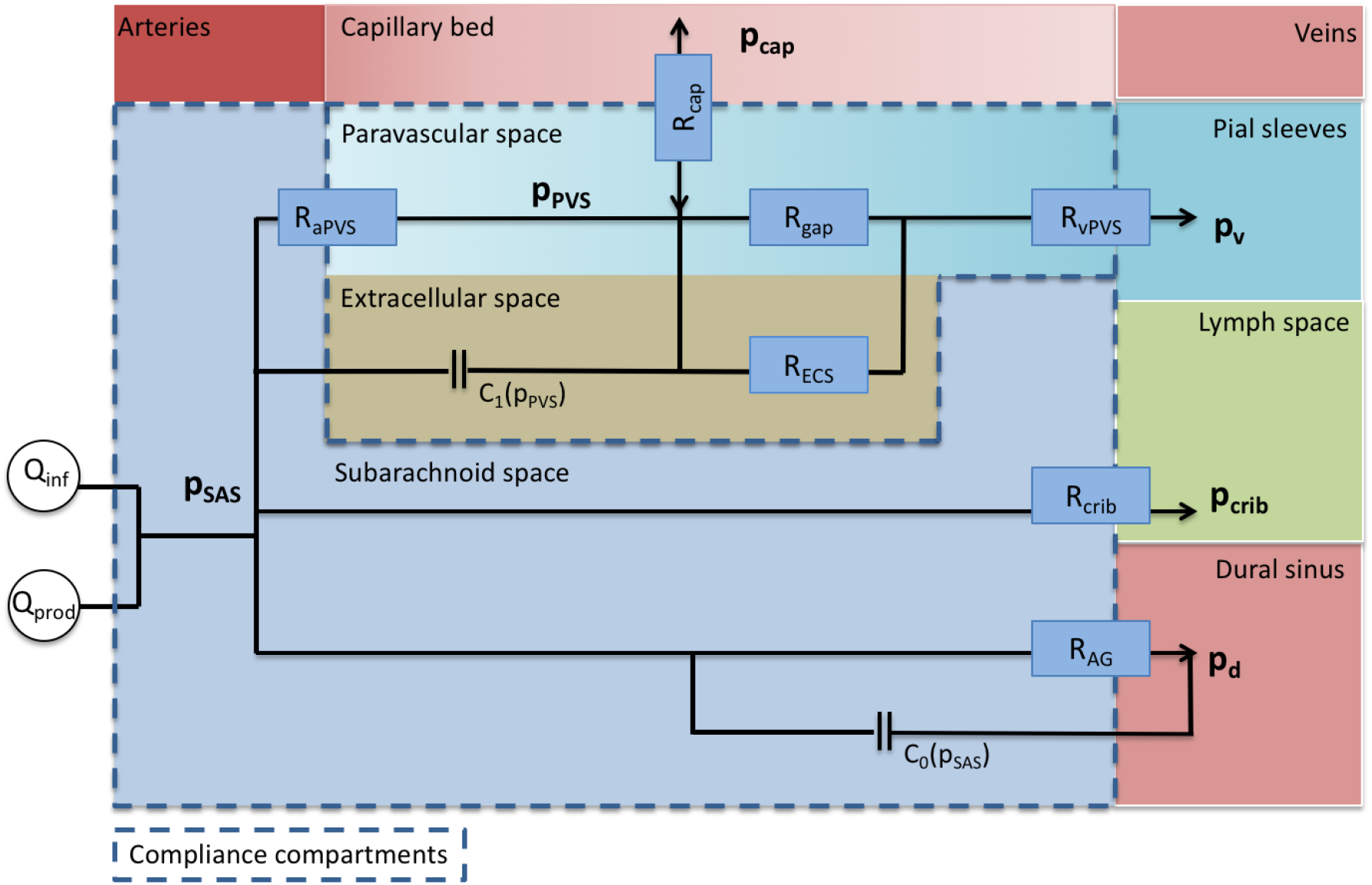
Schematic model description. The model relates the unknown *p*_0_ = *p*_SAS_ and *p*_1_ = *p*_PVS_ and three main exit pathways. CSF formed by production (Q_prod_) and infused fluid (Q_inf_) enter the SAS from the left. The first outflow route is via the arachnoid granulations (AG) where CSF is absorbed by the dural sinuses. The second route is via the cribriform plate (crib), where CSF is absorbed by extracranial lymphatic vessels. In the third outflow route, CSF enters the arterial paravascular spaces (aPVS), from which it may be absorbed by the capillaries (cap) and leave the system via venous blood. Alternatively, the fluid continues along gaps surrounding the capillaries (gaps) or enter the extracellular space (ECS), before entering the venous paravascular spaces (vPVS), where the fluid is assumed to follow PVS along pial veins, via sleeves around the veins, and finally absorbed by the extracranial lymph system. The SAS is considered as one pressure dependent compliance compartment (*C*_0_), and the PVS/ECS represent a second compliance compartment (*C*_1_).

**Figure 2:**
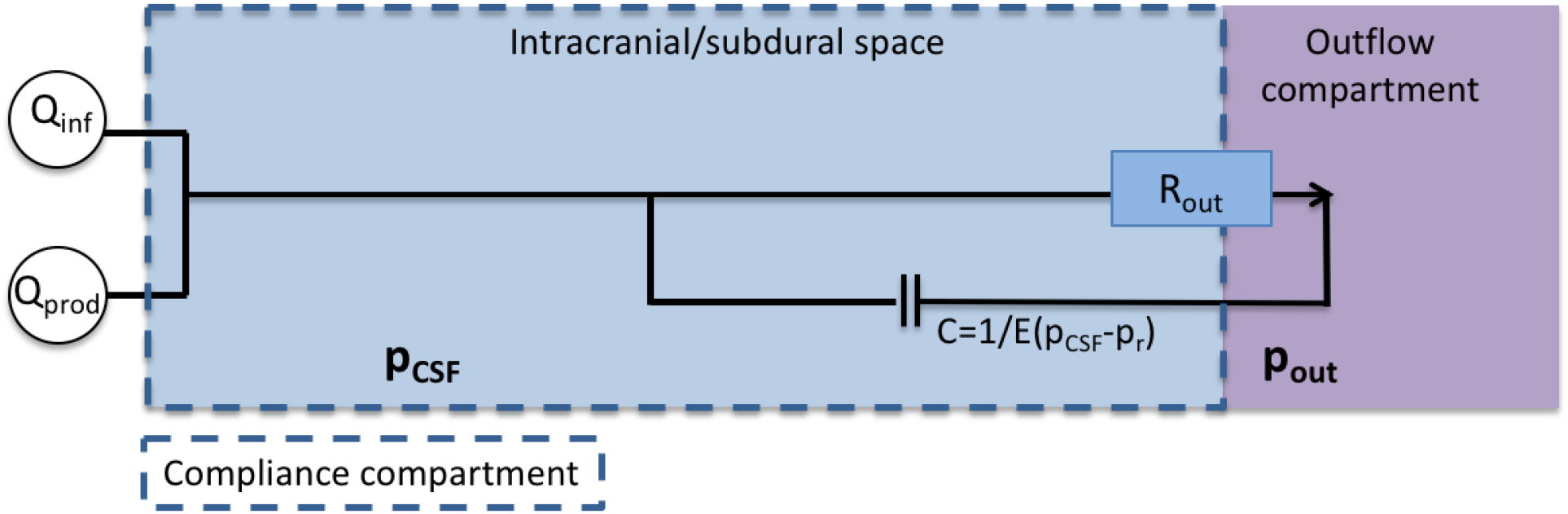
The reference model, an established model for analysis of clinical infusion tests, in which CSF flows out through the arachnoid granulations, with the dural sinus as the outflow compartment and *p*_out_ = *p*_d_. The intracranial resting pressure (ICP_*r*_) is assumed to be related to the unknown *p*_out_, *Q*_prod_, and *R*_out_ by *p*_out_ = ICP_*r*_ – *Q*_prod_*R*_out_.

### Characteristics of infusion resistances and compliance

Infusion tests in humans provide crucial insights and basic characteristics for models of CSF dynamics, including validated pressure ranges. A linear relationship between steady state pressure elevation and infusion rate, i.e. the assumption of a pressure-independent *R*_out_, has been shown to be valid from baseline ICP (*p*_base_) up to *p*_base_+12 mmHg [2]. At higher pressure increases from baseline (>15 mmHg increase), this assumption does not hold. Further, the craniospinal compliance is typically assumed to be inversely dependent on the ICP. This assumption is for most subjects valid from about baseline ICP (approximately 11 mmHg) and higher. For lower ICPs, compliance should be modelled as constant [52, 51]. These insights are reflected by the model proposed here. We assume the inverse compliance model for ICP above baseline pressure and a constant compliance for lower ICP, while the pressure independent property of R_out_ sets the upper ICP pressure validity limit to approximately 23-26 mmHg.

### Mathematical model of CSF pressure dynamics under infusion

We here present a system of two ordinary differential equations (ODEs) describing CSF pressure in the SAS (*p*_0_) and in the PVS/ECS (*p*_1_). For a schematic overview of the model compartments and routes, see Figure 1. Our model extends previous models [14,16] for CSF pressure, flow and compliance within the intracranial compartment, by including the PVS/ECS compartment and additional outflow pathways. To account for flow into the PVS from both capillaries and the SAS, the PVS/ECS is modeled as a pressure compartment similar to the SAS. This compartment represents the PVS from the arteriole to precapillary segments at the end of the paraarterial tree as modeled by Faghih & Sharp [19], and is modeled as in direct contact with the ECS.

The ODE system reads as: find *p*_0_ = *p*_0_(*t*) and *p*_1_ = *p*_1_(*t*) for *t* ≥ 0 such that

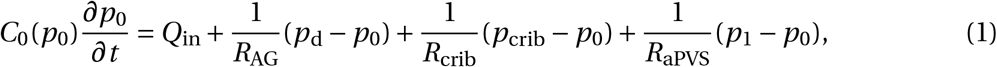

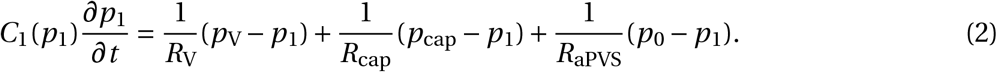

Here, *C_i_* is the compliance of the associated compartment for *i* = 0,1. The (given) counter pressures for the AG, cribriform plate, pial sleeves around veins, and capillaries are denoted by *p*_d_, *p*_crib_, *p*_V_ and *p*_cap_, respectively. The respective resistances are denoted by *R*_AG_, *R*_crib_, *R*_V_ and *R*_cap_. *R*_aPVS_ is the resistance to flow between the SAS (*p*_0_) and PVS (*p*_1_) compartments through paravascular spaces, and *Q*_in_ is the sum of CSF production and fluid infusion. Possible capillary filtration is not included in *Q*_in_, but rather as a separate term in Equation (2). The total resistance of flow *R*_V_ from the arterial PVS through either the capillary gaps or the ECS and finally through the venous PVS is calculated by

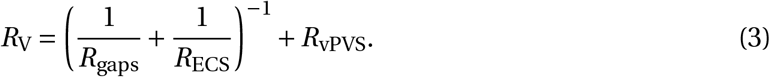

Furthermore, the total resistance as seen from the SAS through the paravascular pathway (glymphatic pathway) can be computed as

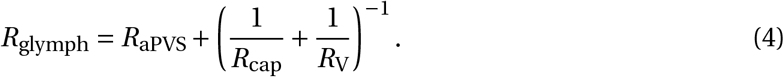

### Model parameters

This section describes the model parameters including estimation of resistances between the different pressure compartments. These parameters are summarized in Table 1.

**Table 1:** Model 1 default parameters (corresponding to full model) cf. Figure 1 and Equations (1)–(2).

| Parameter | Symbol | Value | Unit | Reference |
| --- | --- | --- | --- | --- |
| Dural sinus pressure | $p_d$ | 8.8 | mmHg | Eq. 16 |
| Cervical lymph pressure | $p_{crib}$ | 2 | mmHg | [60] |
| Pial sleeves pressure | $p_v$ | 8.8 | mmHg | model assumption |
| Capillary pressure | $p_{cap}$ | 25 | mmHg | [7] |
| Reference pressure | $p_r$ | 9 | mmHg | [32] |
| Baseline pressure | $p_{base}$ | 11 | mmHg | [2] |
| AGs resistance | $R_{AG}$ | 10.81 | mmHg/(mL/min) | [22] |
| Paraarterial resistance | $R_{aPVS}$ | 1.14 | mmHg/(mL/min) | [19] |
| Paravenous resistance | $R_v$ | 22.96 | mmHg/(mL/min) | Eq. (9) |
| Cribriform plate resistance | $R_{crib}$ | 67 | mmHg/(mL/min) | Eq. (3) |
| Capillary wall resistance | $R_{cap}$ | 125.31 | mmHg/(mL/min) | Eq. (15) |
| Subarachnoid space elastance | $E_0$ | 0.2 | $\text{mL}^{-1}$ | [14] |
| Paravascular/ECS elastance | $E_1$ | $10E_0$ | $\text{mL}^{-1}$ | App.A |
| CSF production rate | $Q_{prod}$ | 0.33 | mL/min | [17, 24, 45, 61] |
| Infusion rate | $Q_{inf}$ | 1.5 | mL/min | [14] |

#### Compliances

The intracranial compliance function is often assumed to take the form [43, 4]:

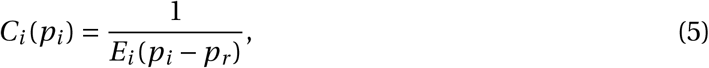

where *E_i_* is known as the elastance coefficient, and *p_r_* is a reference pressure.

For the SAS compartment, we set *E*_0_ = 0.2 mL^-1^ in accordance with reported values [14]. While the physical interpretation of the reference pressure is less immediate [14], we used the value *p_r_* = 9 mmHg as reported by Jacobsson et al. [32]. Furthermore, for *p*_0_ < *p*_base_, where *p*_base_ ≈ 11 mmHg, the assumption of a linear relationship between pressure and volume changes [32], implies a constant *C*_0_ at low pressures. At *p*_0_ < *p*_base_, we thus set the compliance to the baseline compliance in the system.

For the PVS compartment, we assume that the relationship (5) between the PVS pressure *p*_1_ and compliance *C*_1_ holds, but the corresponding parameter *E*_1_ is less known. In a one-compartment model, pressure-volume relationships can be obtained experimentally by infusion of fluid while monitoring the pressure increase. Until a steady state is reached, tissue and possibly other elastic compartments in direct contact with the modeled compartment are displaced due to the increase in pressure. As the SAS and PVS are in direct contact with each other, it is not clear how to distinguish the one from the other in terms of the compliance. Conceptually, it is however clear that the two compliances may be distinct as the SAS space is directly exposed whereas the PVS is exposed through a myriad of narrow channels penetrating deep into the brain.

To determine the elastance parameter *E*_1_ for the PVS compartment, we therefore performed a sensitivity test, aiming for a steady state solution within approximately 15-20 minutes. The results from the sensitivity test are shown in the appendix. With *E*_1_ = *E*_0_, the steady state for a constant infusion test at 1.5 mL/min was not reached until 30–35 minutes. The time to steady state decreased with increasing *E*_1_, and if *E*_1_ = 10*E*_0_, steady state was reached in approximately 15 minutes. The system behaviour did not substantially change further for higher values of *E*_0_. We therefore set *E*_1_ = 10*E*_0_. Similarly as for the SAS, we used the baseline compliance of the system for pressures lower than *p*_base_.

In summary, for *i* = 0,1, we define the compartmental compliance functions

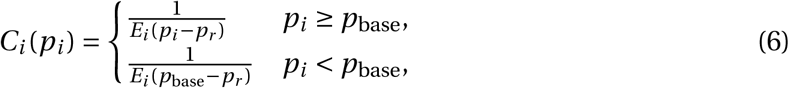

with *E*_0_ = 0.2 mL^−1^, *E*_1_ = 10*E*_0_ and *p_r_* = 9 mmHg.

#### CSF production

In medical textbooks, CSF production rate is normally reported at roughly 500 mL/day [24], which corresponds to 0.35 mL/min. Some studies report a production of around 0.4 mL/min [61, 17], while production rates as low as 0.19 mL/min have been observed in healthy elderly [45]. In this work, we used a production rate of *Q*_prod_ = 0.33 mL/min.

#### Resistance to flow along the microcirculation (*R*_gaps_)

Resistance to flow is given by the ratio of pressure drop (Δ*p*) to the flow rate in the given geometry

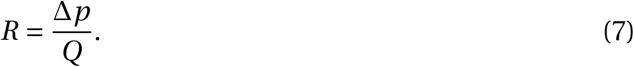

In this work, we will assume the PVS to form a circular annulus around the vessel wall. The flow rate in an annular section is given by [57]:

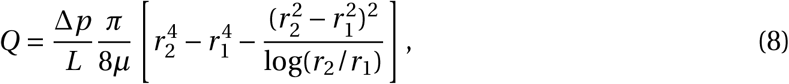

where *μ* is fluid viscosity, *r*_1_ and *r*_2_ are inner and outer radius of the annulus, respectively, and Δ*p* is the pressure drop over the section of length *L*. By combining Equations (7) and (8), the flow resistance in each PVS can be computed as

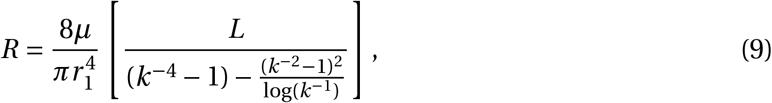

where 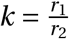 is the ratio between inner and outer radius.

The paravascular resistance model proposed by Faghih and Sharp [19] ended at the precapillary level, where precapillary diameters were 12.2-12.5 *μ*m. In a 1D network of the microvasculature, Payne & El-Bouri [48] reported blood vessel radius and length based on the modeling framework by Boas et al. [6]. The 1D network started at the arteriole level with a vessel radius of 12 *μ*m and branched out to 64 capillaries each with a radius of 4 *μ*m. At the final venule, the radius was 15 *μ*m. We assumed this tree to form the microcirculation and added small gaps of 100 nm [19] around the vessels of varying size allowing for CSF flow along the microvasculature.

The 1D network model [48] consists of a single tree branching from the first generation of arteriole (precapillaries), splitting at each generation until reaching the middle capillary level (generation 7). At this level, the tree consists of 64 capillaries only. After the capillary branch, the tree joins back together to the last venule (postcapillary), such that the total tree consists of 13 generations. The total number of capillaries in the brain is estimated to be around 100 billion [47]. We therefore assumed the brain to have *N* = 1.5625 billion trees, with 64 capillaries in each tree, each identical to the one used by Payne & El-Bouri [48].

The branching tree can be viewed as a combination of parallel and series circuits. To compute the total flow resistance *R*_gap_ in the gaps around the microcirculation, we computed the resistance of each generation and added these together to form the cumulative resistance for a single tree, *R_i_*:

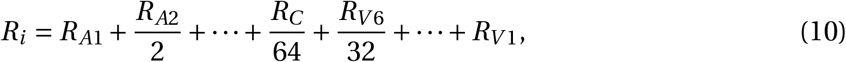

where *R_Ai_* denotes the resistance of the i’th paraarteriolar generation, *R_Vi_* denotes the resistance of the i’th paravenular generation, and *R_C_* denotes the resistance of the paracapillary gaps. Assuming a parallel configuration of the networks, the total resistance of the gaps around the microcirculation is given by dividing the resistance of a single tree by the number of trees *N*:

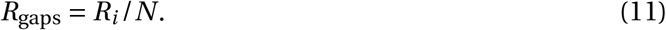

#### Resistance to flow in the ECS (*R*_ECS_)

Estimating the resistance posed by the ECS is a challenge as the ECS is a porous medium in 3D permeated by an almost space-filling network of vessels. Our approach here is to estimate the resistance in terms of a thin disk where the thickness of the disk represents a typical distance between an arteriole and a venule whereas the diameter of the disk represents the size of a human brain. The mean distance from arteriole to venule has been estimated to be *L* = 280 *μ*m in rhesus monkeys [1] and is here used as a lower estimate in humans. Furthermore, we estimate the largest cross sectional area of the human brain forming a disk of radius 7 cm, *A* = *π*0.07^2^ m^2^ ≈ 0.015 m^2^. Finally, the permeability of the ECS has been computed to be *κ* = 10 – 20 nm^2^ in rat neuropil [29]. With these assumptions, we can estimate the resistance using the following relations. First note that the permeability relates pressure and flow by Darcy’s law in a homogeneous medium [11]:

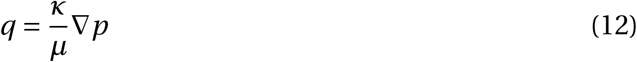

where *q* is Darcy velocity, *μ* is the dynamic viscosity of the fluid, and ∇*p* is the pressure gradient driving flow. Assuming a linear change in pressure across the thickness *L* of the domain (i.e. 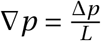), the flow rate is given by

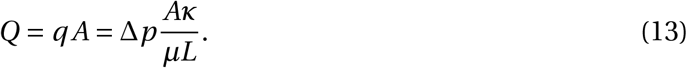

Equations (13) and (7) can be combined to relate permeability to resistance by

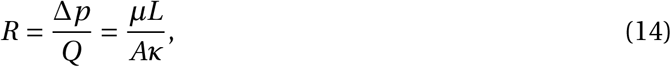

which was used to calculate the total resistance of the ECS.

#### Resistance and pressure for outflow into dural sinus or lymphatics

In AG tissue, the hydraulic conductivity, *L_p_* has been estimated experimentally at 4.52 *μ*L/min per mmHg/cm^2^ (5.66 × 10^-9^ m/(Pa s)) [23], and later adjusted to 92.49 *μ*L/min per mmHg/cm^2^ (1.16 × 10^-7^ m/(Pa s)) with serum free media [28]. The resistance *R* relates to the hydraulic conductivity *L_p_* by

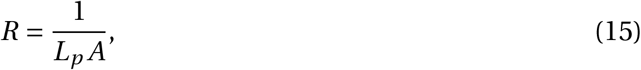

where A is the surface area. An AG surface area of 1 cm^2^ [22] yields *R_AG_* = 221.24 mmHg/(mL/min) without and *R_AG_* = 10.81 with serum free media. The value 10.81 mmHg/(mL/min) seems the more reasonable as median total resistance in healthy elderly has been reported to be 8.6 mmHg/mL/min [40]. Based on the expected results from clinical infusion tests, with *R*_out_ = 8.6 mmHg [40], a resting ICP of 11.6 mmHg [40] and a production rate of 0.33 mL/min, we set the central venous pressure to be 8.8 mmHg according to

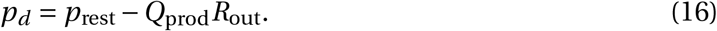

For the AG counter pressure, we thus set *p*_d_ = 8.8 mmHg.

The outflow pressure after the paravenous spaces was assumed comparable to the AG counter pressure at *p_V_* = 8.8 mmHg. Both absorption sites are located in the same anatomical region and both are assumed to be separated from the CSF compartment. Note that this value does not represent the ICP (≈ 10-12 mmHg), nor the lymphatic pressure at the end of the clearance route (2 mmHg). We assume that there are high resistive ducts generating these additional pressure drops.

#### Resistance and pressure for outflow via the cribriform plate

The CSF from the SAS may be transported through the cribriform plate into extracranial lymphatic vessels [60]. The resistance of the cribriform plate has been calculated via infusion tests with and without blockage of the cribriform plate [59]. Assuming an estimated outflow resistance *R*_before_ before blockage and *R*_after_ after blockage, the cribriform plate resistance *R*_crib_ can be estimated by:

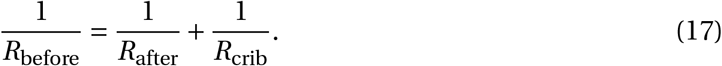

Equation (17) yields a resistance of 67 mmHg/(mL/min).

Cervical lymphatic pressure has been shown (in sheep) to rise only slightly with an increase in ICP [60]. At ICP = 7.4 mmHg, the cervical lymphatic pressure was 1.9 mmHg, for ICP = 22 mmHg, it rose to 2.6 mmHg, and for ICP = 37 mmHg, it rose to 4.4 mmHg. In other words, a constant cribriform plate pressure is a good approximation for the expected ICP values in our model (up to 23 – 26 mmHg). In our model, *p*_crib_ was set to 2 mmHg.

#### Resistance and pressure related to filtration/absorption across capillaries

The total hydraulic (or vascular) conductivity *L_p_* of the blood vessel walls throughout the brain has been reported to be approximately 10^-13^ (m/Pa s) [63]. Assuming a surface-to-volume ratio (S/V) of 10 000 m^2^/m^3^ [63, 67] and that the brain has a volume of about 1 dm^3^ yields a total vascular surface area of A = 10 m^3^. Combining these values with (15), the flow resistance over the blood vessel walls, *R*_Cap_, can be estimated as 125.31 mmHg/(mL/min) (10^12^ Pa s/m^3^).

In peripheral tissue fluid flow across the capillary wall is described by Starling’s law. Fluid is filtrated out on the arterial side of the capillary bed and reabsorbed on the venous side. The difference between filtrated and absorbed fluid is the net flow. Starling’s law has been applied in mathematical models of fluid exchange in the brain [63, 8, 37]. However, both [26] and [9] claim that Starling’s law do not hold for brain tissue due to the low ion permeability of the blood-brain barrier. Fluid exchange across this barrier is still a combination of hydrostatic and osmotic forces [9, 27]. In addition, water may be cotransported along with glucose and oxygen, independent of pressure gradients [39]. The processes involved are complex and not very well understood, therefore, we desregard osmotic effects. The mean capillary pressure was assumed to be 25 mmHg [7].

### Model variations

To investigate the sensitivity of the predicted intracranial pressures and associated flow between compartments with respect to different parameter regimes, we consider a set of model variations. Each variation is labelled and described below, and summarized in Table 2.

**Table 2:** Overview of parameters and modifications for models 0–9 used in the study. All other parameters, cf. Table 1, were kept constant. The resistances *R* are given in mmHg/(mL/min) and pressures *p* in mmHg.

| Model | $R_{AG}$ | $R_{crib}$ | $R_{glymph}$ | $R_{aPVS}$ | $R_{cap}$ | $R_V$ | $p_V$ |
| --- | --- | --- | --- | --- | --- | --- | --- |
| 0 | 8.6 | ∞ | ∞ | ∞ | ∞ | ∞ | 8.8 |
| 1 | 10.81 | 67.0 | 20.54 | 1.14 | 125.31 | 22.96 | 8.8 |
| 2 | 10.81 | ∞ | ∞ | ∞ | ∞ | ∞ | 8.8 |
| 3 | 10.81 | 67.0 | ∞ | ∞ | ∞ | ∞ | 8.8 |
| 4 | ∞ | 67.0 | 20.54 | 1.14 | 125.31 | 22.96 | 8.8 |
| 5 | 21.62 | 67.0 | 20.54 | 1.14 | 125.31 | 22.96 | 8.8 |
| 6 | 10.81 | 67.0 | 0.004 | $1.43 \times 10^{-3}$ | 125.31 | $2.6 \times 10^{-3}$ | 8.8 |
| 7 | 10.81 | 67.0 | 20.54 | 1.14 | 125.31 | 22.96 | 2.0 |
| 8 | 10.81 | 67.0 | 20.54 | 1.14 | 125.31 | 22.96 | $p_0(t)$ |
| 9 | 10.81 | 67.0 | 24.10 | 1.14 | ∞ | 22.96 | 8.8 |

#### Reference model (model 0)

As a reference model, we used the median value of *R*_out_ = 8.6 mmHg/(mL/min) as reported by Malm et al. [40] in a cohort of healthy subjects. In simulations this was accomplished by assuming AG to be the only outflow route, thus setting *R*_AG_ = 8.6 mmHg. All other resistances *R* were set to ∞, i.e., 1/*R* = 0.

#### Full model (model 1)

The model described in Figure 1 and by Equations 1 and 2 is referred to as the full model, or model 1. In this model, all exit pathways considered were assumed possible. The models (2–9) below are described as modifications to the full model.

#### AG as the only outflow route (model 2)

In this modification of the model, we assumed that all flow occurs through the AG, similarly as for the reference model. However, in this modification, we set *R*_AG_ = 10.81 mmHg/(mL/min), the same value as was used in all models except the clinical reference model (model 0).

#### Glymphatic pathway eliminated (model 3)

The glymphatic circulation and its role in the net clearance of CSF out of the intracranial compartment is disputed [18]. We tested the effect of eliminating glymphatic function in our model by letting all flow exit through the cribriform plate or the AG.

#### AG pathway eliminated (model 4)

This model represents a complete dysfunction in the AG, effectively eliminating the AG as a pathway. We tested whether other outflow routes could compensate, and to what extent ICP would be increased.

#### Increased AG resistance (model 5)

Experimentally, the AG outflow resistance has been found to vary by a factor 20, depending on the fluid properties. We therefore also tested an increased resistance of 21.62 mmHg (corresponding to twice the value of the full model) to test the model sensitivity with respect to this parameter. This variation also tested whether other outflow routes could compensate for an increase in *R*_AG_, in contrast to a complete elimination as in model 4.

#### Extended capillary gaps (model 6)

Mestre et al. [46] demonstrate PVS collapse after fixation. This observation entails that a gap size of 100 nm at the precapillary level, as used by Faghih and Sharp [19] and measured after fixation, may be an underestimation. If we rather assume a linear change in area ratio between cross sectional PVS and lumen (going from 1.26 to 0.13 [58] over 13 generations), resistance is vastly reduced. For the different vessel radii reported by Payne & El-Bouri [48], we set 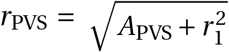, and computed the resistance in Equation (9) with 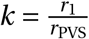.

#### Decreased glymphatic outflow pressure (model 7)

To allow for a potential pressure difference between the inlet and outlet of the intraparenchymal compartments (PVS and ECS), we have introduced an additional compartment (identified with hypothetical pial sleeves) at the venous end of the PVS. We assumed a high pressure (8.8 mmHg) in this compartment compared to the lymph pressure (2 mmHg), but a low pressure compared to e.g. the baseline ICP (11 mmHg). We therefore examined decreasing the paravenous outflow pressure (*p_V_*) to the cervical lymph pressure (2 mmHg) to account for uncertainties in this parameter.

#### Circular glymphatics (model 8)

In this model, which can be viewed as a converse of model 7, we let ISF return directly to the SAS from paravenous spaces, setting *p_V_* = *p*_0_ in Equation (2). In addition, in Equation (1), we add the term 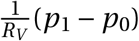 to account for the possibility of fluid to leave and enter the SAS also along the paravenous space. The net flow into and out of the brain will thus be by filtration and absorption over the capillary wall. We also assumed extended capillary gaps as for model 6 and thus reduced PVS resistance compared to the full model.

#### Capillary filtration eliminated (model 9)

Even though capillaries constantly filtrate water [9], the net flow rate over the capillary wall, if any [26, 39], is hard to quantify. We therefore also tested the effect of eliminating flow over the capillary wall from our model.

### Computation of steady state values

In steady state, the left hand side of Equation (1) is zero and independent of compliance. We calculated the steady state solution of the CSF pressure in the SAS as a function of the infusion rate *Q*_inf_. The function take the form

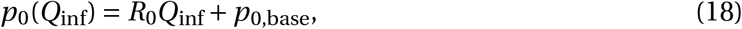

In a clinical infusion test, *R*_out_ is measured in the SAS, and thus corresponds to our estimation of *R*_0_. *p*_0,base_ is the ICP in the SAS at baseline ICP. For each model we report both *R*_0_ and *p*_0,base_ such that ICP can be computed for any arbitrary infusion rate. In addition, we calculate the pressures and corresponding flows at rest and at peak ICP resulting from an infusion test with *Q*_inf_ = 1.5 mL/min.

### PVS velocity estimation

Following [19], we modeled the paraarterial tree as stemming from three branches of the arterial tree. The paraarterial model started at generations 18,16 and 16 after the middle cerebral artery, anterior cerebral artery, and second posterior cerebral artery with diameters *d*_0_ = 100.16, *d*_1_ = 97.42 and *d*_2_ = 97.42 *μ*m (and radii *r*_0_, *r*_1_ and *r*_2_), respectively [19]. Following the assumption that arterial PVS area is *c* = 1.26 times that of the lumen area [58], the total PVS cross-sectional area at the starting branch is:

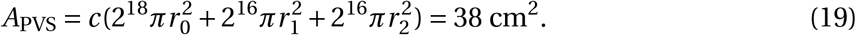

We can thus calculate the average velocity at the base of the arterial PVS by

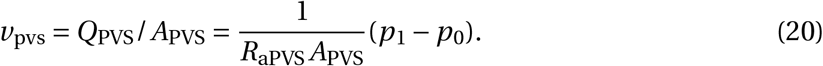

for ICP pressure *p*_0_ and PVS pressure *p*_1_.

### Numerical methods and setup

The system (1)-(2) of ODEs was solved in Python using the Scipy [34] (version 1.1.0) ODE solver, odeint. At each time step, the solver recognizes characteristics of the linear system and is adaptive with respect to solver method (Adams or BDF), order, and time step. As initial conditions, we set *p*_0_(0) = *p*_1_(0) = 8 mmHg, and we assumed a CSF production *Q*_prod_ = 0.33 mL/min (see 1). The equations were first solved to reach a steady state (t = 60 min) before adding the infusion fluid *Q*_inf_ = 1.5 mL/min for 30 minutes. Thus in Equation (1), we set

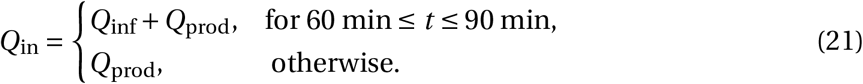

We note that the given *Q*_prod_ does not include the possible capillary filtration resulting from Equation (2).

## Results

In the three following subsections, results for the full model (model 1) is presented along with comparison to the reference model. In the fourth subsection results from modifications to the full model (models 2-9) are presented.

### Flow resistances

#### Capillary gaps

For the single 1D network [48], the resistance was computed to be

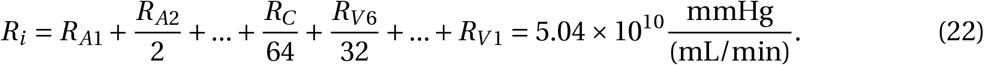

Assuming the parallel configuration of the networks, the total resistance of the annular gaps around the microcirculation is found by dividing by the number of trees:

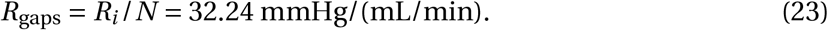

This resistance is far too high for paravenous flow to re-enter the SAS, as any considerable flow would require local pressure differences of several mmHg in the SAS. For instance, a flow of 0.13 mL/min, representative of interstitial fluid perfusion in humans [19], would require a pressure drop of 4.19 mmHg along the capillary gaps. The maximal estimated transmantle pressure gradient of 0.03 mmHg [64] would drive less than 0.9 *μ*L/min through the glymphatic circulation.

### ECS

To estimate a lower bound for the resistance in the ECS, we used the highest permeability and the lower bound of the distance between arteriole and venule. Combining Equations (13) and (7) with *κ* = 20 nm^2^, L = 280 *μ*m, A= 0.015 m^2^ and viscosity *μ* = 0.7× 10^-3^ Pas, we arrive at

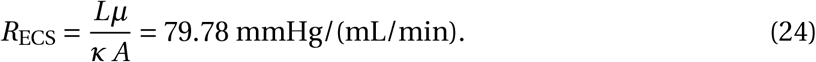

The computed resistance in the ECS is relatively high, but not more than a factor 5 higher than an upper estimate of the normal total outflow resistance [40].

### Flow distribution between compartments

Figure 3 shows the results from a standard infusion test with a production rate of 0.33 mL/min and constant infusion at a rate of 1.5 mL/min. Baseline ICP was 11.07 mmHg, and peak ICP was 20.68 mmHg, as compared with 11.66 and 24.56 in the reference model (model 0). The outflow SAS resistance (as defined by Equation (18) was *R*_0_ = 6.41 mmHg/(mL/min). The full model thus had lower resistance to outflow from the SAS than the reference model, but was still within the variability of *R*_out_ as seen in healthy subjects [40]. For this reason ICP was higher in the reference model, and flow through (the only outflow route) the arachnoid granulations were greater in the reference model.

**Figure 3:**
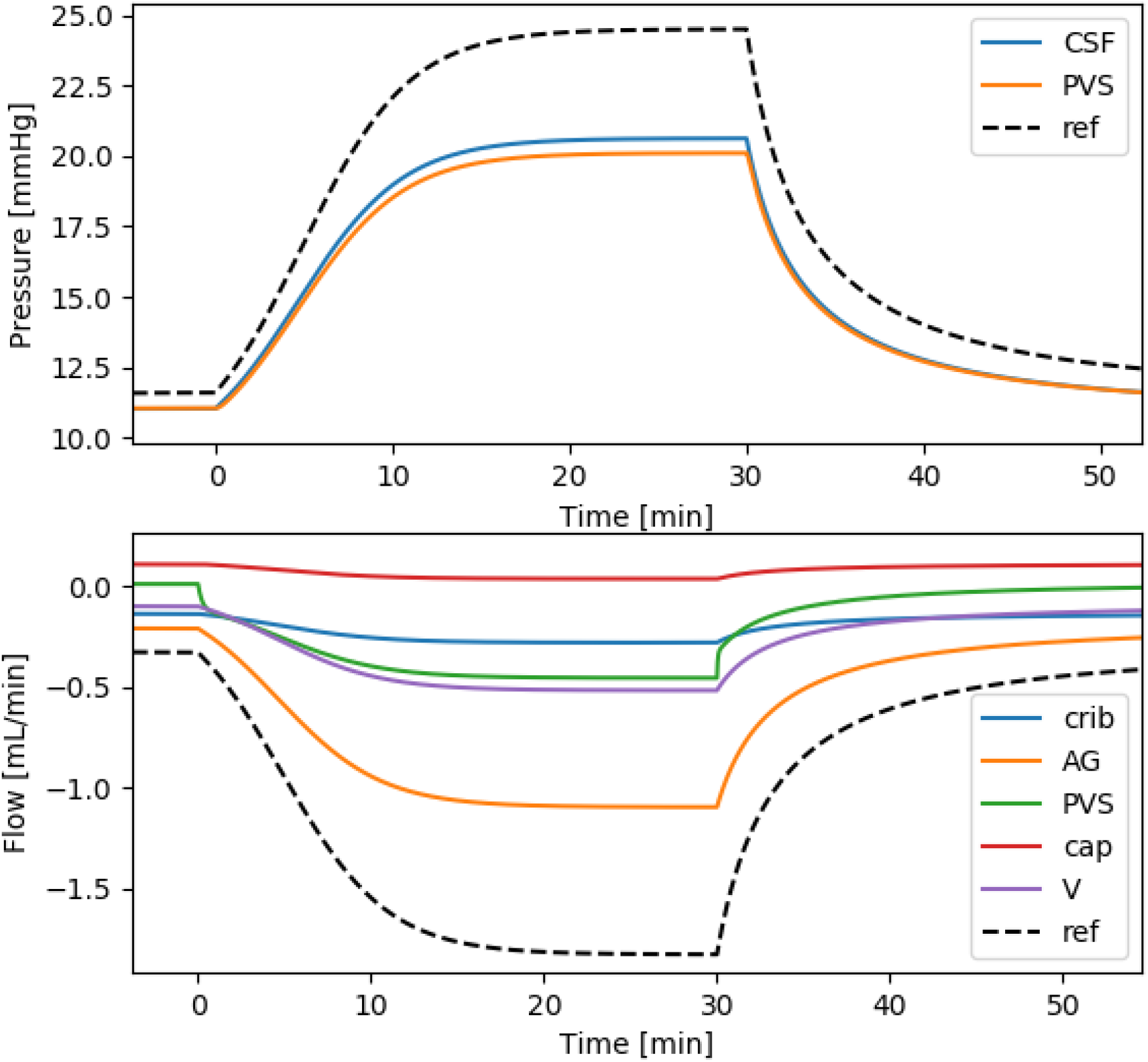
CSF pressure and outflow during a standard infusion test. The arachnoid granulations dominated outflow both under baseline and (elevated) plateau ICP, with 60.0% and 60.1% of the fluid leaving through the granulations. The flow to pial sleeves (*p_V_*) and eventually to cervical lymph (purple curve) is greatest at plateau, and equals the paravascular flow (green) minus capillary filtration (red) when the system is in steady state. The cribriform plate plays a much less prominent role at plateau than at baseline ICP (40.0% vs 15.3%). At baseline, paravascular flow is almost stagnant: the amount of fluid entering the SAS from the PVS is only 2.9% of the total outflow. Importantly, PVS flow reverses during the infusion test and accounts for 24.6% of the clearance out from the SAS (and into the PVS) at plateau ICP.

At baseline ICP, flow through the AG was 0.21 mL/min. 0.01 mL/min went from the PVS into the SAS, while 0.14 mL/min left the system through the cribriform plate. Net capillary filtration was 0.11 mL/min, leaving the system through the ECS or capillary gaps, through the paravenous spaces, towards the pial sleeves and eventually the lymphatic system. The distribution of clearance from the SAS to each compartment was thus found to be 60.0% to AG, and 40.0% to the cribriform plate. The PVS flow entering the SAS was only 2.9% the value of the total outflow from the system.

At plateau ICP, clearance was also dominated by flow through the AG. However, the PVS play a more prominent role. 1.10 mL/min was cleared through the granulations, while for the PVS and cribriform plate, the flow rate was 0.45 and 0.28 mL/min, respectively. Net capillary filtration was 0.03 mL/min. At plateau ICP the distribution of clearance from the SAS to the different compartments changed, with 60.1% exiting through the granulations, 24.6% into the PVS and 15.3% through the cribriform plate. Our main findings are also illustrated in Figure 4.

**Figure 4:**
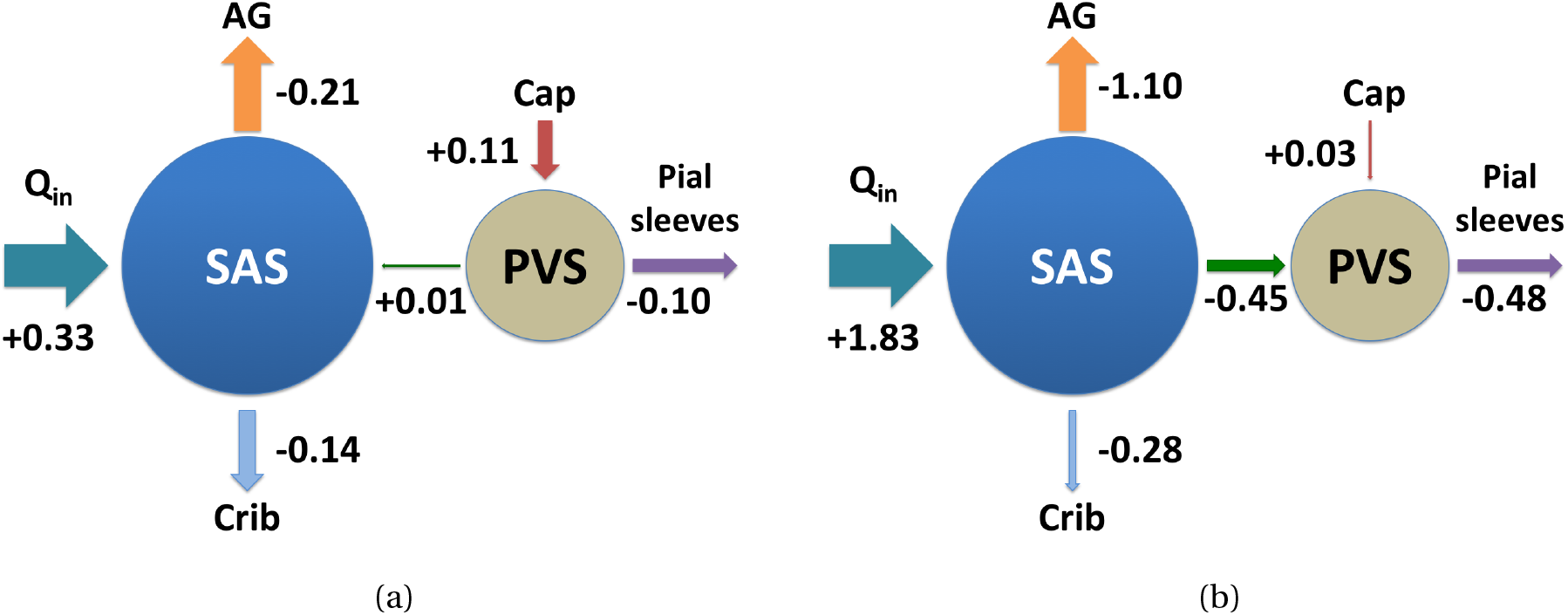
Illustration of the outflow distribution in mL/min (a) before and (b) during infusion. The size of the arrows are proportional to *Q*_in_. Note that the net CSF flow from the PVS to the SAS changes direction and magnitude during infusion.

### PVS velocity and pressure

At baseline, the PVS pressure was 11.08 mmHg as compared to 11.07 mmHg for the ICP (Figure 3). During the infusion, ICP increased faster than the PVS pressure, but the two curves stayed relatively close throughout the infusion test. The pressure difference increased at plateau ICP, where the PVS pressure was 20.16 mmHg, while ICP was 20.68 mmHg.

Using Equation (20), with *A*_PVS_ = 38 cm^2^ and flow rate as given in Table 3, the resulting PVS velocities were *v*_PVS_ = 0.04 *μ*m/s into the SAS at baseline and *v*_PVS_ = 1.97 *μ*m/s into PVS after infusion.

**Table 3:** Effect of modifications to steady state results of the model on ICP and flow. Results are reported as max/min where max is the value at plateau ICP (shown within parenthesis), while min is the value at baseline ICP. Pressure is given in mmHg, flow in mL/min and resistance in mmHg/(mL/min). Negative values indicate flow out from the SAS and into the relevant compartment. Similarly, for capillary flow negative values indicate flow from PVS into capillaries.

| Mod | ICP ($p_0$) | AG flow | PVS flow | Crib flow | Cap flow | $R_0$ |
| --- | --- | --- | --- | --- | --- | --- |
| **0** | **11.66 (24.56)** | **-0.33 (-1.83)** | **n/a** | **n/a** | **n/a** | **8.60** |
| 1 | 11.07 (20.68) | -0.21 (-1.10) | 0.01 (-0.45) | -0.14 (-0.28) | 0.11 (0.03) | 6.41 |
| 2 | 12.39 (28.60) | -0.33 (-1.83) | n/a | n/a | n/a | 10.81 |
| 3 | 10.95 (24.91) | -0.20 (-1.49) | n/a | -0.13 (-0.34) | 0.11 (0.00) | 9.31 |
| 4 | 14.34 (37.92) | n/a | -0.15 (-1.29) | -0.18 (-0.54) | 0.09 (-0.10) | 15.72 |
| 5 | 12.01 (25.67) | -0.15 (-0.78) | -0.03 (-0.70) | -0.15 (-0.35) | 0.10 (-0.00) | 9.10 |
| 6 | 8.82 (8.83) | -0.00 (-0.00) | -0.23 (-1.73) | -0.10 (-0.10) | 0.13 (0.13) | 0.00 |
| 7 | 9.27 (18.88) | -0.04 (-0.93) | -0.18 (-0.65) | -0.11 (-0.25) | 0.13 (0.05) | 6.41 |
| 8 | 11.92 (24.92) | -0.29 (-1.49) | 0.10 (0.00) | -0.15 (-0.34) | 0.10 (0.00) | 8.66 |
| 9 | 10.35 (20.43) | -0.14 (-1.07) | -0.06 (-0.48) | -0.12 (-0.27) | n/a | 6.71 |

### Model variations

The models stayed within the assumption of a plateau pressure below 26 mmHg (as discussed in the Methods section), with the exception of model 2 and model 4. The exact output values from these latter two models could thus be affected by the assumption of a pressure independent *R*_out_. *R*_0_, as defined in Equation (18), denotes the total outflow resistance from the SAS compartment. *p*_0,base_ is defined as baseline ICP, and is given as the first number under the ICP column. Instead of reporting the PVS pressure for all models, we rather report the PVS flow.

The PVS pressure can be deduced from the PVS flow via Equation (7) as *p*_1_ = *p*_0_ + *Q*_aPVS_*R*_aPVS_. Here, *Q*_aPVS_ is the PVS flow as reported in Table 3, *p*_0_ is the pressure in the SAS and *R*_aPVS_ is the paravascular resistance.

#### Reference model (model 0)

The reference model behaved as expected, with a pressure increase from 11.66 to 24.56 during constant infusion at 1.5 mL/min. The plateau ICP in the SAS was at the transition phase for when the assumption of a pressure independent *R*_out_ is valid (p > 23-26 mmHg).

#### Full model (model 1)

The results from the full model was described more in detail in the previous three sections. However, we note that the resistance to outflow from the SAS was reduced in the full model compared to the reference model (Table 3, model 1 vs. model 0).

#### Only AG clearance (model 2)

Outflow only through the AG using resistance as estimated experimentally [28] resulted in higher ICP than with the reference model. However, the increase from the reference model was less than 1 mmHg at baseline, and approximately 4 mmHg at plateau ICP (Table 3). Compared to the full model, the ICP increase at plateau was the most dramatic change for model 2 (≈8 mmHg higher than the full model).

#### Glymphatic pathway eliminated (model 3)

With the glymphatic pathway eliminated, ICP was practically unchanged at baseline compared to the full model (a 0.12 mmHg decrease). At plateau ICP however, elimination of the glymphatic pathway increased ICP to 24.91 mmHg, more than 4 mmHg greater than full model (Table 3). Outflow was greatest through the AG. For baseline ICP, the relative flow through AG and the cribriform plate stayed approximately the same as in the full model. AG was by far the most important outflow route at plateau ICP (1.49 vs 0.34 mL/min for the AG versus the cribriform plate).

#### AG pathway eliminated (model 4)

Compared to the full model, total elimination of the AG as outflow route increased ICP substantially to 14.34 mmHg and 37.92 mmHg at baseline and plateau ICP, respectively (Table 3). At plateau, ICP was greater than the capillary pressure, a critical threshold value, where capillaries absorb rather than filtrate water. At baseline, capillaries filtrate 0.09 mL/min, while at plateau 0.10 mL/min is absorbed. Also in model 4, PVS flow and flow through the cribriform plate is similar at baseline (0.15 vs 0.18 mL/min), but during infusion, flow is dominated by PVS flow (1.29 vs 0.54 mL/min). Model 4 had the highest PVS flow (except model 6) at 1.29 mL/min out from the SAS. This flow rate corresponds in a PVS velocity of 5.66 μm/sec.

#### Increased AG resistance (model 5)

Double AG resistance caused a decrease in AG flow from 0.21 to 0.15 mL/min during baseline compared to the full model (Table 3). Similarly as for model 4, PVS flow and flow through the cribriform plate was similar during baseline ICP (0.08 vs 0.10 mL/min), and the relative importance of PVS increased at plateau (0.75 vs 0.31 mL/min). At plateau, ICP was almost equal to the capillary pressure, and approximately zero net filtration/absorption occurs. Increased AG resistance gave ICP and *R*_0_ closer to the reference model (model 0) compared to model 1.

#### Extended capillary gaps (model 6)

In model 6, the calculated resistance for the extended capillary gaps was found to be 0.9 × 10^-3^ mmHg/(mL/min). With the assumptions described for model 6 in the methods section, the smallest capillary gaps were computed to be 1.2 *μ*m, a result seemingly more in agreement with figures by Pizzo et al. [49]. Following the steps from Faghih & Sharp [19], we calculated the resistance in the paraarterial tree, without the assumption of smaller gaps on the final generation in their model, to be 1.48 × 10^-3^ mmHg/(mL/min). This calculation seems to be in good agreement with their presented figure of cumulative resistance in the paraarterial tree. With these adjustments, most CSF immediately left the system through PVS, capillary gaps and eventually to lymph nodes through the paravenous route. There was a small, steady flow (0.10 mL/min) out through the cribriform plate due to the relatively large pressure difference between the cribriform counter pressure and the ICP. For the AG, the counter pressure is approximately equal to the ICP, and no flow occurs.

#### Decreased glymphatic outflow pressure (model 7)

In view of the conjectural nature of the paravenous route to the lymphatics, a decrease in the counter pressure after the paravenous route was assumed in model 7. The outflow pressure was set at 2.0 mmHg. ICP was decreased compared to the full model, down to 9.27 at baseline and 18.88 at plateau ICP (Table 3). In this case, outflow was predominantly into the PVS at baseline and through AG at plateau.

#### Circular glympahtics (model 8)

If the paravenous route connects back to the SAS, no directional glymphatic circulation is possible. In other words, in our model, paraarterial and venous flow always had the same direction: either both from the SAS into the PVS or both from the PVS to SAS. The PVS flow reported in Table (3) for this model is thus the sum of paraarterial and paravenous flow entering the SAS compartment. In this model, flow was directed out from the PVS as capillary pressure stayed higher or equal to the ICP and PVS pressure. At baseline ICP, the capillary filtration rate (and thus, net flow to the SAS from PVS) was 0.10 mL/min, while at plateau pressure, the net filtration/absorption rate was 0 mL/min. The flow rate along arterial PVS was 0.07 mL/min, and the venous PVS flow was 0.03 mL/min at baseline ICP. From the SAS, the main outflow route was the AG, where the flow rate was twice the flow rate through the cribriform plate at baseline, and more than four times greater at plateau.

#### Capillary filtration eliminated (model 9)

Compared to the full model, a balanced filtration/reabsorption across the capillary wall altered the results slightly at baseline, but results were nearly identical at plateau ICP. At baseline, flow through the AG and crib were almost identical (0.14 vs 0.12 mL/min, Table 3), and PVS flow was always directed from the SAS into PVS. However, the paravascular flow rate was small at baseline (0.06 mL/min) and increased by a factor of 8 at plateau ICP (0.48 mL/min).

## Discussion

In this work, we computed CSF resistance, pressure and outflow both during resting state and during infusion. The mathematical model extended the traditional infusion modeling with pathways related to the glymphatic system. In addition, we calculated PVS pressure, flow and average velocity in both states. All models except model 6, which involved significant flow in the PVS at the capillary level, gave reasonable ICP and ICP increase from a constant infusion of fluid at a rate of 1.5 mL/min. The best match between the traditional reference model and the proposed model were achieved by increasing the resistance of the AG by a factor two (model 5).

Our estimates of resistances of capillary gaps and ECS were on the same order as resistances to other outflow routes such as the cribriform plate and arachnoid granulations as measured by others [23, 28, 60]. However, the resistance in the capillary gaps or through the ECS was about 30 times greater than the resistance in the paraarterial tree used in our model, the latter based on estimations by Faghih & Sharp [19]. In particular, this increase in resistance renders flow through arterial PVS continuing to venous PVS and a return to the CSF even less plausible than what was discussed by Faghih & Sharp [19]. In our case, a reasonable flow rate would require a pressure drop of 4.19 mmHg. Such flow must rely on local ICP differences in the SAS, and would be expected to induce flow directly along the SAS rather than through the possibly high-resistant glymphatic system. In addition, pressure gradients in the SAS are likely less than 3 mmHg/m [44, 55], and the transmantle pressure difference has been estimated to be no more than 0.03 mmHg [64]. Glymphatic circulation driven by local differences in pulsatile pressure of several mmHg also seems implausible, as the ICP wave is almost identical in time everywhere in the brain [13], with a maximal estimated pulsatile difference of approximately 0.2 mmHg [68]

Ma et al. [38] suggest that outflow through the cribriform plate dominates, but only when the total outflow from the SAS is large. We also found the relative distribution of flow to the different outflow pathways to be affected by infusion, but there are important differences. Under no conditions did we find the flow through the cribriform plate to dominate flow to the other outflow routes as was suggested in their study. With the full model, at baseline ICP, AG flow was 50% greater than flow through the cribriform plate, and PVS flow nearly absent. AG flow was greater or equal to flow through the cribriform plate in all cases in all models except for model 6. The relative distribution to different outflow routes have been assessed under both awake and anaesthetized conditions. While we did not address this question, there is conflicting evidence whether PVS flow increase or decrease during sleep or anesthesia [38, 21, 69]. To what extent pressure, outflow resistances, or CSF production cause changes between different states is not well understood.

Net capillary flow in our model was mostly directed from the capillaries to the PVS. Thus, under normal conditions, the capillaries functioned as a site of CSF production. In models where pressure exceeded 25 mmHg, the capillaries functioned as a site of absorption. It is interesting to note that this pressure threshold is exactly at the value for which R_out_ becomes pressure dependent as measured experimentally [32]. Capillary flow rates were relatively small, up to 0.13 mL/min. Capillaries functioning as a route of absorption and filtration of CSF is in line with the Bulat-Klarica-Orešković hypothesis [9]. However, it should be noted that passage of substances such as proteins and electrolytes is difficult over the blood-brain barrier as compared to water [20]. The routes of CSF/water clearance (CSF is 99% water [9]) does not necessarily align perfectly with the routes of clearance of other substances from the brain. We finally note that estimations of capillary resistance one order of magnitude lower than we used has been estimated by Koch [36].

The average PVS velocity in the full model was 0.04 out from the PVS at baseline and 1.97 μm/s into the PVS at plateau ICP. The maximal recorded PVS velocity (not including model 6) was computed to be 5.66 μm/s in model 4. In the experimental studies, several investigators used an infusion rate of 2 μL/min in rodents [30, 31, 46], which was shown to increase ICP by 2.5 mmHg [31], a substantial increase, but less than in our model of a human with an infusion rate of 1.5 mL/min. In addition, Bedussi et al. [5], used a much lower infusion rate of 0.34 μL/min, only resulting in a pressure rise of 0.1 mmHg. Still, the net PVS velocity of 17 μm/s found with this low increase in ICP is almost identical to the typical flow speed of 18.7 μm/s found by Mestre et al. [46] at the infusion rate of 2 μL/min, suggesting elevated ICP is not the sole reason for PVS flow. These findings do not support our finding of reversal of the PVS flow direction during infusion. However, up to this point, experimental studies able to measure paravascular flow velocities have been limited to rodents, and as such are not directly comparable to our model.

According to Tithof et al. [66], PVS are not concentric cylinders, but rather form ellipses around vessels to minimize resistance. This geometrical change along all vessels may decrease resistance by a factor of 2-3 [66], and thus likely to increase PVS velocities by a similar factor in our model. In addition, peak velocity in a concentric cylinder is double that of the mean velocity, which possibly may increase our velocity estimates of up to 5.66 μm/s in the models by another factor of approximately two.

The current study gives new insight to the relation between ICP and CSF clearance. This is not only highly relevant to the glymphatic theory, but also to understand pathological conditions related to increased ICP. Patients with increased ICP often exhibit clinical visual symptoms, which are also typical in astronauts suffering from Spaceflight Associated Neuro-ocular Syndrome (SANS)[15].

### Limitations

Resistance parameters in our model were taken from many different species and types of experiments. Ideally, resistance in each outflow pathway could be measured experimentally by blocking one outflow pathway, and measure the resulting increase in pressure. To our knowledge, this has not been done in humans and it is likely challenging both from a technical and ethical perspective.

The introduction of a hypothetical compartment connecting the venous PVS to the dural lymphatics give rise to a hydrostatic gradient large enough to drive flow through the PVS. However, it should be noted that the original glymphatic theory as described by Iliff et al [30] also suggest a route from the paravenous spaces to the SAS.

Our model did not include the effect of cardiac or respiratory pulsatility neither in the arterial, venous or CSF compartments, which all has been proposed to drive glymphatic clearance [31, 46, 5]. In particular, the cardiac pulsation on the arterial side seems related to the PVS pulsatile movement as shown in Mestre et al. [46], but modeling attempts deem it unlikely that arterial wall movements alone drive a net flow of sufficient magnitude for clearance of fluid [3, 54].

We did not consider spatial variation, but rather assumed a compartment model, where pressure was a function of time only. ICP has been shown to be nearly identical in space [13], whilst the blood flow pulse propagation has a given directionality [35, 53].

Capillary filtration is regulated by osmotic gradients [9,27] and cotransporting proteins [39], which were not considered in our study. With the simplifications made in our study regarding spatial variation, inclusion of Starling forces would correspond to a reduction in the capillary hydrostatic pressure. Absorption may thus still occur even if the hydrostatic pressure in capillaries exceeds the pressure in surrounding tissue or PVS. In addition, the capillary pressure was assumed constant in time, also in models where ICP increased to, or even higher than the capillary pressure. Whether the capillary pressure always stays above ICP regardless of ICP increase is not well known. In model 9, we ensured that alterations in the relative distribution from an infusion test was changed also when capillary filtration was removed from the model.

## Conclusions

According to our models of CSF clearance, outflow predominantly occurred through the arachnoid granulations, although the relative distribution to each outflow route were dependent on ICP. Paravascular flow occurred from the PVS to the SAS at baseline ICP and was reversed at plateau ICP. We conclude that ICP increase is an important factor to address when determining the pathways of injected substances in the SAS.

## Author’s contributions

VV, AE, MER, KAM, KHS developed the mathematical model and designed the study. VV and KHS performed the parameter estimations. VV performed the simulations. VV and KHS created the figures. VV prepared the first manuscript draft. VV, AE, MER, KAM and KHS edited the manuscript. All authors approved the final manuscript.

## Acknowledgements

We thank Dr. Eric Schmidt, Centre Hospitalier Universitaire de Toulouse France, for discussions and useful insight on infusion testing.

## A Elastance sensitivity

As discussed in section about model parameters we performed a sensitivity analysis for the elastance of the PVS/ECS compartment (E_1_). The result is shown in Figure 5.

**Figure 5:**
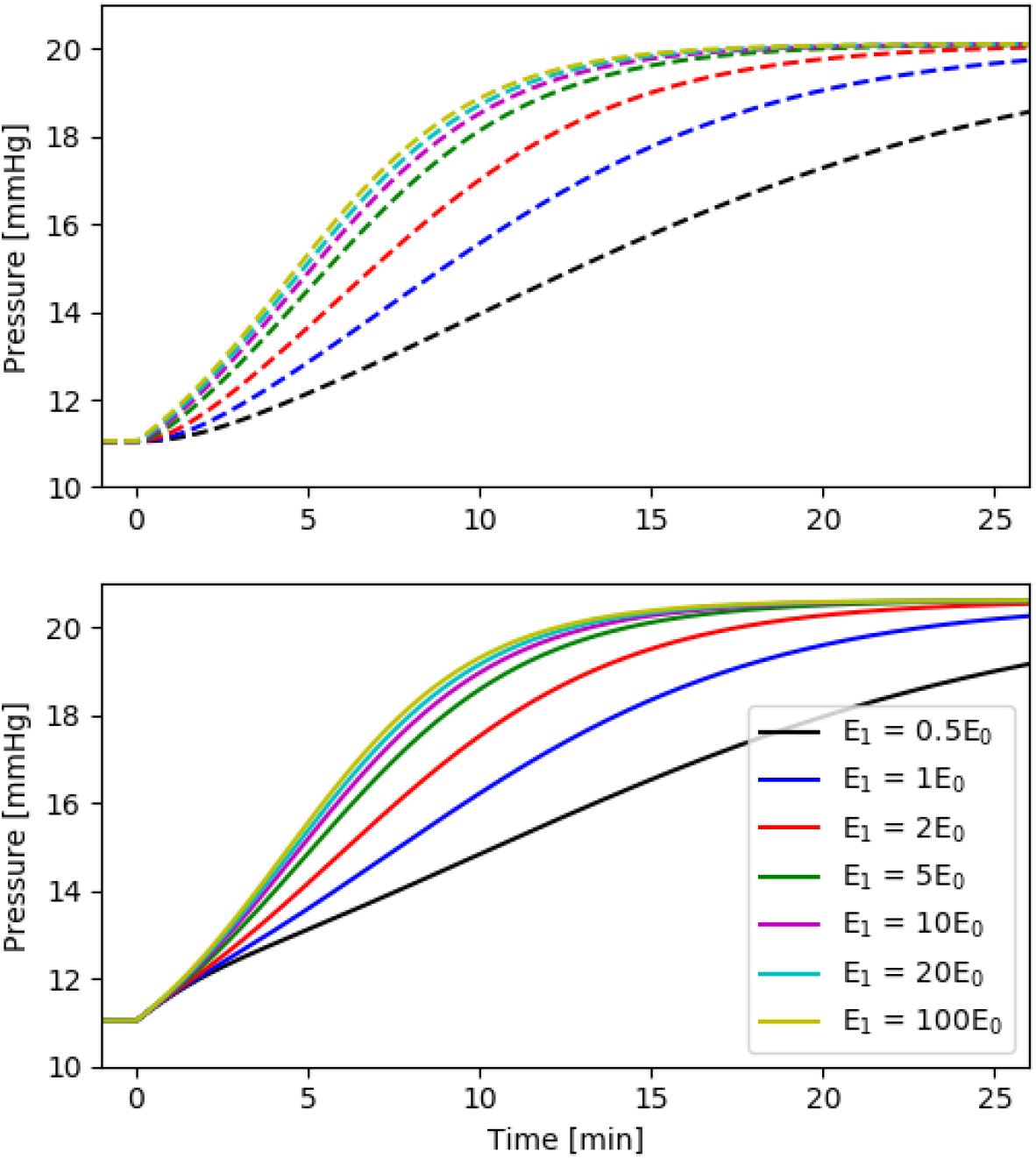
Elastance sensitivity for the evolution of *p*_PVS_ (top) and *p*_SAS_ (bottom). Simulation setup as for the full model (model 1), but for different values of E_0_. For the SAS elastance we used E_1_ = 0.2 mL^-1^ as before.

